# Turing pattern prediction in three-dimensional domains: the role of initial conditions and growth

**DOI:** 10.1101/2023.03.29.534782

**Authors:** Soha Ben Tahar, Jose J Muñoz, Sandra J Shefelbine, Ester Comellas

## Abstract

Reaction-diffusion systems have been widely used to model pattern formation in biological systems. However, the emergence of Turing patterns in three-dimensional (3D) domains remains relatively unexplored. A few studies on this topic have shown that extending pattern formation from 2D to 3D is not straightforward. Linear stability analysis, which is commonly used to associate admissible wave modes with predicted patterns in 1D and 2D, has yet to be applied in 3D. We have used this approach, together with finite element modelling of a Turing system with Schnakenberg kinetics, to investigate the effects of initial conditions and growing domains on the competition between admissible modes in 3D Turing pattern emergence. Our results reveal that non-random initial conditions on the activator play a stronger role than those of the inhibitor. We also observe a path dependency of the evolving pattern within a growing domain. Our findings shed new light on the mechanisms ensuring reliable pattern formation in 3D domains and have important implications for the development of more robust models of morphogen patterning in developmental processes.

## 1 Introduction

The process by which organisms develop their shape and form during embryonic development has fascinated scientists for centuries. Pattern formation is a crucial stage in morphogenesis, as it leads to the emergence of structures that later support function. Alan Turing proposed a mechanism to explain how cell signalling can generate self-organising patterns, which drive cell differentiation and organisation into specific tissues and structures. In his influential paper [1], Turing modelled the behaviour of chemical signals, which he termed morphogens, through a reaction-diffusion system in which a spatial pattern emerges as a result of diffusion-driven instabilities. The formation of Turing patterns depends on the delicate balance between diffusion and reaction rates, as well as the nonlinear feedback between chemical species.

Turing systems have been extensively used to model patterning in a variety of biological applications [2,3], and the past two decades have brought experimental evidence to support Turing-type computational model predictions [4–9]. Most studies to date have focused on one- and two-dimensional models, with only a handful of papers considering reaction-diffusion systems in three-dimensional (3D) domains to explore patterning in development [10–12].

For more than half a century, Turing models have been the focus of research, but questions remain regarding the selection and emergence of specific patterns. Despite a simple and widely applicable mathematical framework, the nonlinearities inherent to the reaction-diffusion system make it challenging to predict pattern evolution. Linear analysis is a common technique to assess the stability of the steady-state system and investigate pattern formation. It has been used to predict patterns in 1D [3] and 2D structures [13–17], albeit with limitations. To account for the effect of the nonlinearities on pattern emergence, more sophisticated mathematical analyses have been used [18–25]. Numerical methods like finite difference and finite element analyses have become the standard tool in the study of Turing pattern formation.

To date, a limited number of computational studies have examined the emergence of Turing patterns in 3D domains [19, 20, 26–33]. The extension from 2D to 3D domains leads to a wider variety of patterns. All these studies have focused on the generation of complex patterns. However, we know from linear stability analysis in 1D and 2D that these patterns are in fact the superposition of simpler patterns (e.g. spheres, cylinders and planes in 3D), which are associated to a specific wave mode. Admissible modes for a specific set of model parameters are obtained from the linear analysis of the governing equations, while the combination of the admissible modes leading to the final pattern is influenced by initial conditions [13, 21, 29, 30, 34–36].

Through a combination of computational modelling and linear stability analysis, we have investigated how the modes determine the emergence of Turing patterns in 3D domains as well as the effect that initial conditions have on them. A better understanding of mode superposition in 3D will lead to insights on how more complex patterns form. Morphogen expression observed in developmental processes like embryonic axis specification or limb formation typically results in relatively simple patterns (e.g. gradients or ellipsoids). With this type of application in mind, we have also explored how growth may affect the evolution of the pattern as the domain grows.

## 2 Theoretical background

### 2.1 A reaction-diffusion model with Schnakenberg kinetics

The general dimensionless form of a two-component Turing system is given by the reactiondiffusion equations

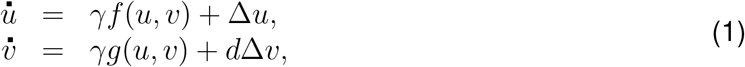

where *u*(***x***, *t*) and *v*(***x***, *t*) are the species concentrations. Their time derivatives are indicated by a superimposed dot, and 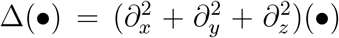 is the standard Laplacian operator. Time and space dependence will be in general omitted for clarity. The term *γ* can have multiple interpretations linked to the domain size and to the relative strength of the reaction terms, and *d* is the ratio of diffusion coefficients of each species. The functions *f* (*u, v*) and *g*(*u, v*) will be here represented by the Schnakenberg kinetics [37], i.e.,

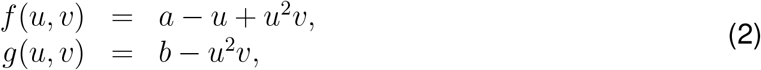

where *a* and *b* are the positive parameters of the model. Hence, the reactant *u* acts as an activator by self-activating itself and inhibiting *v*, while *v* self-inhibits itself and activates *u*, which corresponds to an activator-substrate model.

There are several advantages to using the Schnakenberg general nondimensional form. The first one is that it has a relatively large Turing space in comparison to other models, making the Schnakenberg model more robust [38]. In addition, other well-known systems, such as the Gierer-Meinhardt model [39], can be scaled to take this general form. Another advantage is that the parameters *γ* and *d* have a physical and biological interpretation.

### 2.2 Pattern prediction using linear stability analysis

Linear stability analysis of the Turing system’s steady-state solution is commonly used to study the conditions necessary for the emergence of patterns [17, 25, 40]. The most unstable mode of the resulting eigenvalue problem typically determines the wavelength of the pattern. Thus, this sort of analysis is a useful tool in determining the critical parameter values under which diffusion-driven instability will occur and Turing patterns will form.

The emergence and stability of patterns is analysed by assuming solutions of (1) linearised at ***u***_0_ = (*u*_0_, *v*_0_) = (*a* + *b, b/*(*a* + *b*)^2^) and with the form,

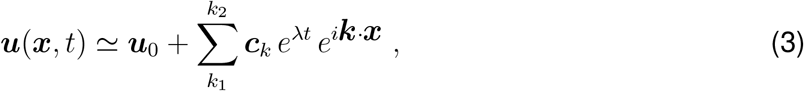

where ***k*** is the wavevector, and the constants ***c***_*k*_ are determined by a Fourier expansion of the initial conditions. The sum in (3) runs on a set of plausible values of *k* = *∥****k****∥*, to be specified below. The stability of the system is determined by the sign of the real part of *λ*, which can be written in terms of *k*. Indeed, after inserting the expression in (3) into the linearisation of (1), the presence of non-trivial solutions requires that *λ* depends on *k*^2^ as

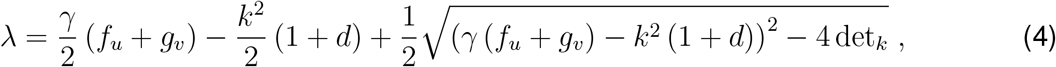

where *f*_*u*_, *f*_*v*_, *g*_*u*_ and *g*_*v*_ are the derivatives of the Schnakenberg kinetics evaluated at (*u*_0_, *v*_0_) and function det_*k*_ is the determinant of the matrix in the linearised system. See Supplementary Information (SI), section S1 for more details.

The relation in (4) is the so-called *dispersion relation*, and allows relating the wavelength *k* with the (positive) real part of *λ*. The emergence of plausible (unstable) oscillatory modes requires that 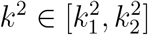, with the values of the interval given by

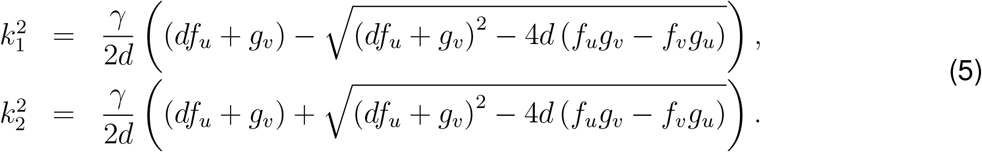

We will focus our study on parallelepipedic domains of dimensions *L*_*x*_ *× L*_*y*_ *× L*_*z*_ and with homogeneous Neumann boundary conditions (BCs). In this case, the wavevectors ***k*** in (3) must take the form

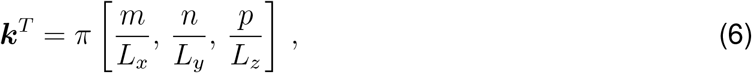

where the integers *m, n* and *p* represent the number of oscillations along the *x, y* and *z* directions, respectively. We have chosen Neumann BCs as they provide a better approximation of the biological environment in morphogenesis than Dirichlet BCs. Note also that in our parallepipedic domain, adding periodic BCs restricts the set of solutions to even integer numbers in (6), while solutions with homogeneous Neumann BCs can take odd and even integers. In view of (6), the condition 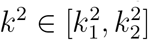 is tantamount to requiring that

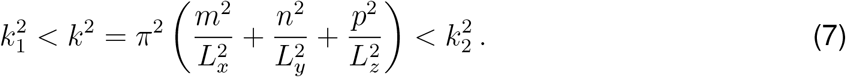

In consequence, there is only a finite and discrete number of integers *{m, n, p}* and values of *k* for which *k*^2^ is within the interval 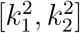. Model parameters and domain dimensions may be selected such that only a single wavelength *k* verifies (7), i.e. there exists a single admissible mode (SI, section S1) and, according to the linear analysis, only this mode will appear in the final pattern. This is referred to as “mode selection” by Murray [3], where the use of linear stability analysis to predict patterns in 1D and 2D domains is explained in detail. We note though that the approximated linearised solution in (3) does not satisfy the nonlinear equations in (1)-(2), and therefore, the steady-state solutions obtained in our simulations cannot be exactly identified with the expression in (3).

When several wavelengths are interacting nonlinearly, the wavelength with the maximum *λ* is expected to grow faster and be the dominant contribution. We denote by (*m, n, p*) the resulting *pure mode* with ***k*** = *π* [*m/L*_*x*_, *n/L*_*y*_, *p/L*_*z*_] and *k*^2^ satisfying the inequalities in (7). When *multiple* modes are interacting, complex patterns may form, which result from the combination of the patterns corresponding to pure modes.

## 3 Numerical simulations

Through computational modelling in 3D domains we examined how the modes predicted by the solution to the linearised system (3) contribute to the final pattern computed with the nonlinear governing equations (1)-(2). We also explored the impact of initial conditions and a growing domain on Turing pattern emergence.

### 3.1 Methods

The governing equations (1)-(2) were discretised in space by applying the finite element method and in time using the implicit mid-point rule. Details of the numerical implementation in Matlab (2022a, The MathWorks Inc.) are provided in SI, section S2. The time step was Δ*t* = 0.1 and simulations ran for as many increments as required until the pattern reached a steady-state solution. Steady-state was considered to be reached at *t*_*n*_ when the difference of nodal values of *u* and *v* where all below 0.5% with respect to values at *t*_*n*_ *-* 10Δ*t*. In random initial conditions we considered a perturbation of up to a 10% variation from a constant value corresponding to ***u***_0_ = (*u*_0_, *v*_0_) = (*a* + *b, b/*(*a* + *b*)^2^). Cubic geometries were meshed with 20*×*20*×*20 hexahedral elements and homogeneous Neumann BCs were applied. The model parameters *a, b* and *d*, and domain sizes *L*_*x*_ *× L*_*y*_ *× L*_*z*_ were adjusted for each simulation. Values are given alongside the results for each case. For simplicity, *γ* = 1 was considered in all simulations, unless stated otherwise.

Figure 1 shows three final patterns obtained from a *pure mode* (*m, n, p*) each. These allow us to unequivocally associate a mode with its graphical representation (or pattern) in 3D, up to half-length translations and symmetries. We targeted the mode by choosing specific boundary conditions, model parameters and domain sizes. To expedite the numerical computation, we considered an initial distribution of the reactant concentrations close to the targeted mode. In the results shown in figure 1, we applied periodic and homogeneous Neumann boundary conditions, which only allow an even number of oscillations, and facilitate the association of the mode to a pattern.

**Figure 1:**
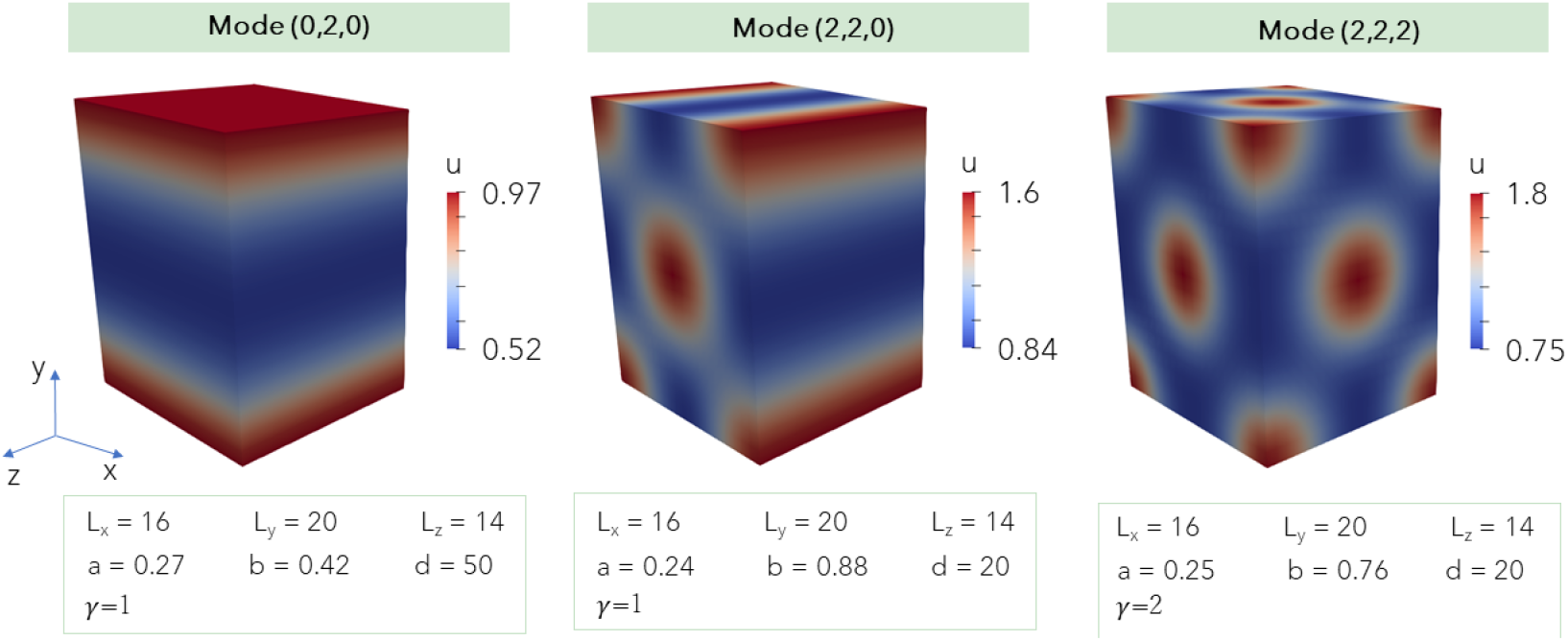
Graphical representation of pure modes in 3D for the reactant u. Applying specific initial conditions, and periodic and homogeneous Neumann boundary conditions allows obtaining a final pattern that corresponds to a pure mode, when selecting a particular geometry and set of model parameters.

Mode (0, 2, 0) corresponds to a full cosine wave in the *y*–direction, which results in a pattern with a plane perpendicular to the *y*–direction (figure 1, left). Mode (2, 2, 0) corresponds to full cosine waves in the *x*– and *y*–directions, which form central and corner cylinders with their axes perpendicular to the *x*–*y* plane (figure 1, centre). Finally, mode (2, 2, 2) is the product of three full cosine waves, one in each direction, which result in a set of split spheres (figure 1, right), with a pattern on each plane similar to the pattern on the *x*–*y* plane of mode (2, 2, 0). We note that in all cases, the central values of *u* are close to the value *u*_0_ = *a* + *b* predicted by the linear analysis.

### 3.2 Mode selection and its influence on the final pattern

We performed several simulations in which we chose the parameters and domain size to select certain admissible modes and, therefore, target specific patterns. We found that the final pattern sometimes corresponds to the mode associated with the largest eigenvalue of the admissible wavenumbers, but not always. Figure 2 shows two illustrative cases: when the final pattern is associated with the largest eigenvalue (C) and when it is not (F). To aid in the identification of the pattern corresponding to the predominant mode, we ran additional simulations (A, B, D, E) in which we tailored the model parameters to obtain only one of the possible eigenvalues within the admissible range of wavenumbers of figure 2C and F. For each simulation (A-F), the positive real part of the dispersion relation (4) is plotted and the wavelengths verifying (7) are indicated on the plot.

**Figure 2:**
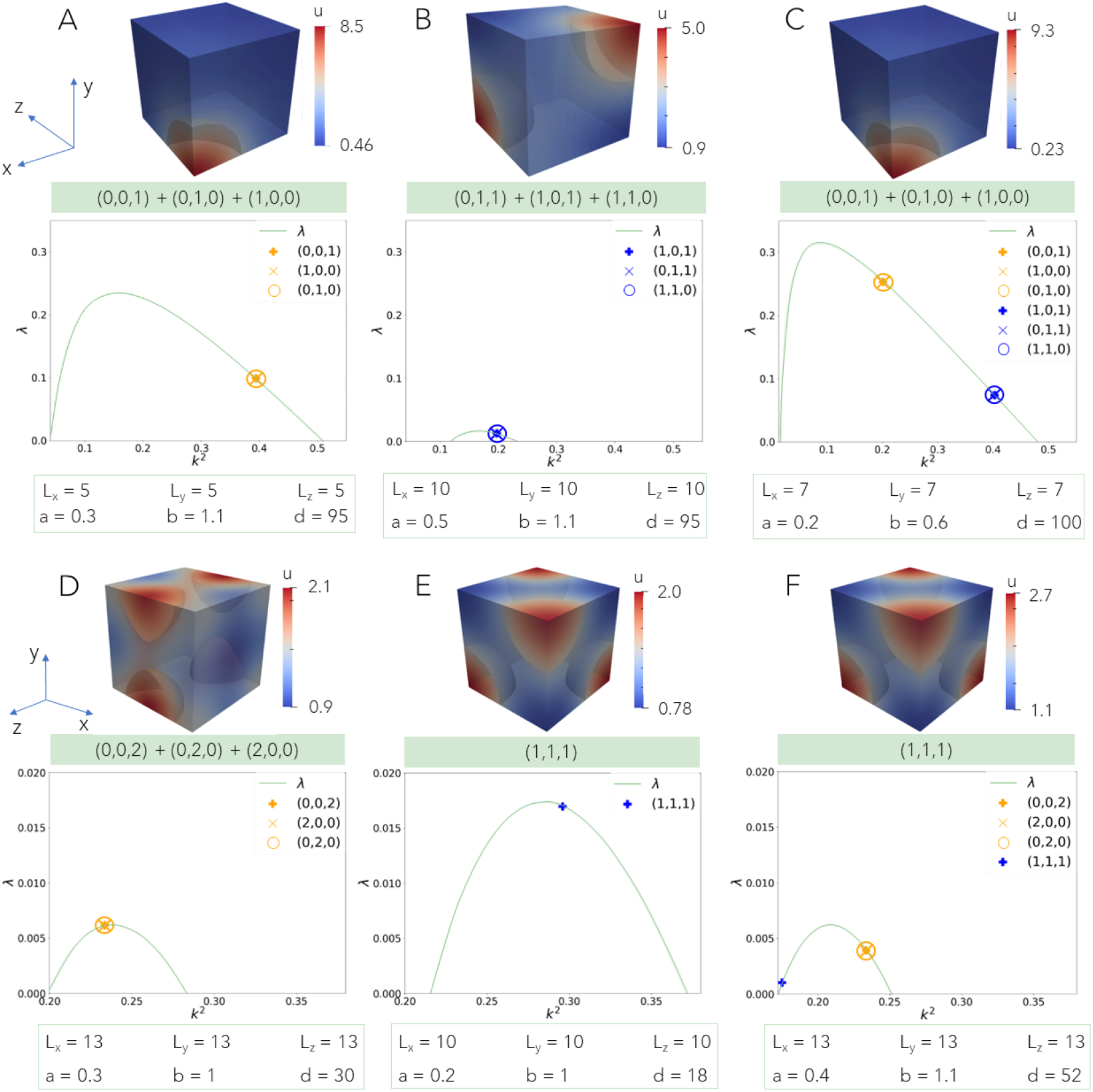
The mode corresponding to the largest eigenvalue is not necessarily the predominant one in the final pattern. For each case (A-F), the plot represents the dispersion relation *λ* vs the wavenumber *k*^2^ obtained from the linear stability analysis (equations (4)-(7)). We can deduce the admissible modes and their corresponding eigenvalue from it. Below each plot, the domain dimensions and model parameters considered are given. On top of the plot, the steady distribution of *u* obtained by finite element analysis is shown. The modes participating in the pattern obtained from the non-linear system are specified below each simulation. Each figure shows patterns associated with corresponding admissible modes. The final pattern in C corresponds to the sum of the modes (0, 0, 1)+(0, 1, 0)+(1, 0, 0), which have the largest eigenvalue. Conversely, the final pattern in F corresponds to mode (1, 1, 1), which has the lowest eigenvalue.

The number of modes interacting depends on the domain dimensions and the model parameters *a, b* and *d*. Most final patterns are a combination of several admissible modes predicted by the linear analysis. The multiplicity of modes with a same eigenvalue is typically known as *mode degeneracy* [26, 29]. For the example in figure 2A, the three modes obtained from the linear analysis are (0, 0, 1), (0, 1, 0) and (1, 0, 0). Each of them separately correspond to a plane perpendicular to the *z*–, *y*– and *x*–directions, respectively. Added together, the final pattern forms an eighth of a sphere in one corner of the cubic domain. We note that this is qualitatively different from mode (1, 1, 1) shown in E and F, as illustrated also in SI, figure S1. Therefore, the final pattern in A results from an equal contribution of the three admissible modes.

Similarly, for the example in figure 2B, the final pattern results from the addition of the three modes obtained through the linear analysis. Modes (0, 1, 1), (1, 0, 1) and (1, 1, 0) are individually associated with two quarter cylinders in opposite corners parallel to the *x*–, *y*– and *z*–planes, respectively. Added together, these modes form an eighth of a sphere in two opposite corners of the cubic domain.

Figure 2C shows that the dominant mode of the final pattern corresponds to the largest eigenvalue (the addition of modes (0, 0, 1), (0, 1, 0) and (1, 0, 0)), similar to the pattern observed in figure 2A. The opposite is true for the example of figure 2F, in which the dominant mode obtained, (1, 1, 1), corresponds to the smallest eigenvalue. This final pattern is identical to the one predicted for the example in figure 2E, for which there is a single admissible mode. Note that this pattern, in contrast to the one in figure 2F, corresponds to what we have termed a pure mode as it does not result from a combination of modes, and corresponds in fact to one eight of mode (2, 2, 2) in figure 1 (right).

Figure 2D shows the pattern associated with the sum of modes (0, 0, 2), (0, 2, 0) and (2, 0, 0). It corresponds to the largest eigenvalue in figure 2F, but for latter parameters and dimensions, it does not seem to contribute to its final pattern. It appears that when eigenvalues are close together, as in this case, the nonlinearities of the reaction-diffusion system may result in a smaller eigenvalue predominating over the largest one, making final pattern prediction based on the linear analysis unreliable. In these cases, the initial conditions become critical in determining the final result. This is explored in the next section.

### 3.3 Impact of the initial conditions on the final pattern

Initial conditions play a critical role during the mode competition described in the previous section. A specific initial condition can favour one mode over the other. Leppä nnen et al. [30] showed that even random initial conditions can favour one type of pattern over another.

To study the impact of initial conditions on the final pattern, we performed a series of simulations considering the same geometry, boundary conditions and model parameters as that of figure 2C (case 1 in figure 3) and 2F (case 2 in figure 3), but starting from different random initial conditions with up to a 10% or 20% variation around the constant values (*u*_0_, *v*_0_).

**Figure 3:**
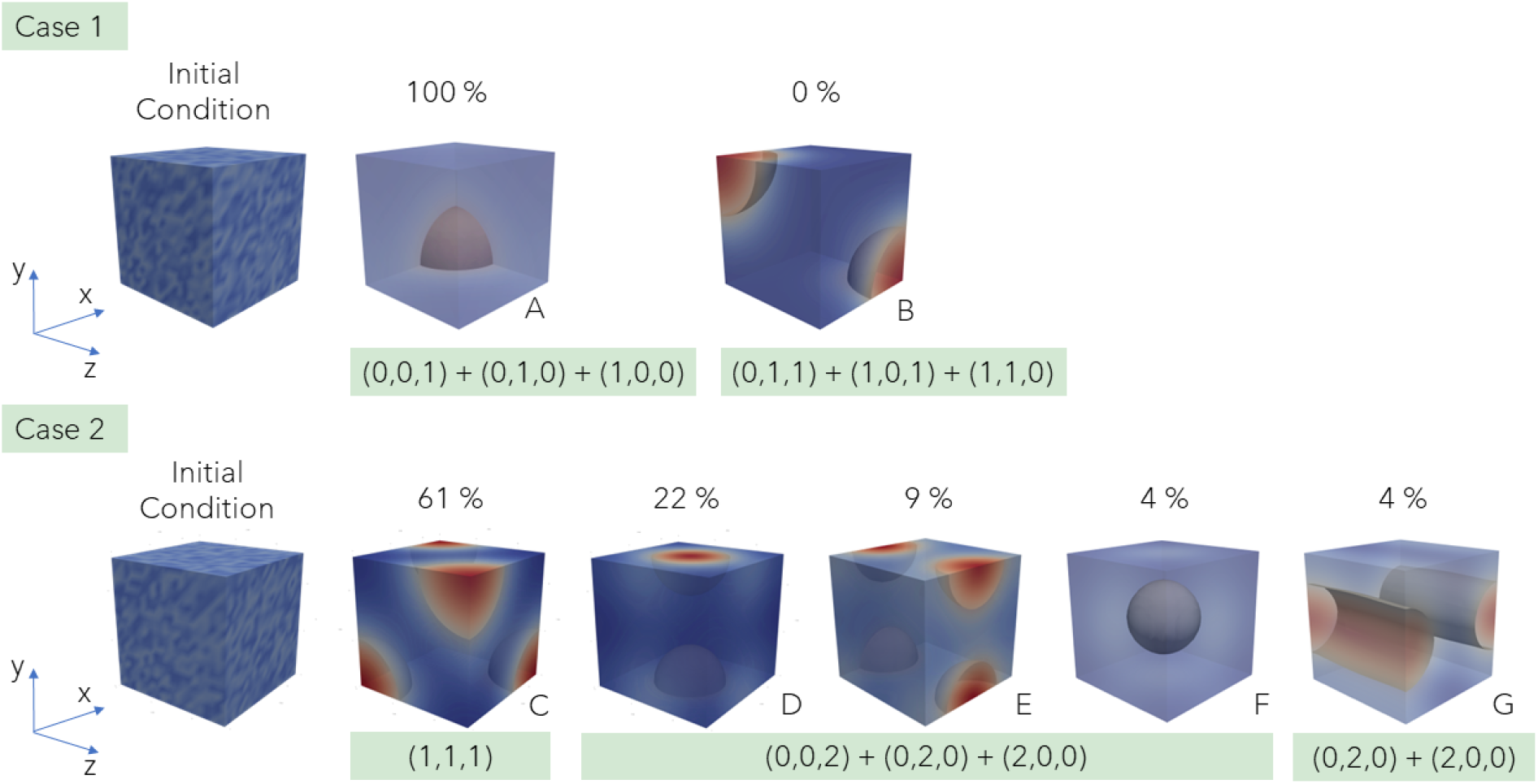
Effect of different random initial conditions on the final pattern of u. The model parameters, boundary conditions and domain size correspond to the ones of figure 2C (case 1) and figure 2F (case 2). The frequency of appearance of each pattern is indicated as a percentage out of the total of 23 simulations performed for each case. The modes that contribute to each final pattern are specified below the patterns. The pattern corresponding to the mode that does not appear in case 1 (B) has been obtained from a different set of model parameters and is shown for illustrative purposes. The highest *u* values are shown in red, and the lowest in blue. While case 1 always resulted in the same final pattern (A), for case 2 different patterns were obtained depending on the randomness of the initial conditions (C-G).

Figure 3 shows that, for particular configurations with the exact same conditions except for different randomness in the initial conditions, different patterns may be obtained. For case 1, we consistently predicted the same final pattern, regardless of the initial conditions (figure 3A). Case 2 was very sensitive to initial conditions (figure 3C-G). The difference between the two cases may be explained by the interval between the eigenvalues (see figures 2C and 2F). When eigenvalues are close to each other, the resulting final pattern is more sensitive to the initial conditions and multiple final patterns may emerge. For case 2, mode (1, 1, 1) appeared the most (14 of 23 simulations). Note that the patterns observed in figure 3D-F are in fact identical if translated half the length of the domain, and correspond to the combination of modes (0, 0, 2), (2, 0, 0) and (0, 2, 0).

This example illustrates how the symmetries of homogeneous Neumann BCs may span the possible patterns arising from the simulation, suggesting that boundary conditions play a strong role during mode competition and the frequency of their appearance. In case 2, the equivalent patterns in figures 3D-F corresponding to combinations of modes with the smallest eigenvalue, were obtained in 61% of the simulations while those corresponding to the largest value were obtained in 39% of the simulations. The pattern in figure 3G corresponds to the sum of modes (2, 0, 0) and (0, 2, 0), and further illustrates how modes may combine in unexpected ways to form different final patterns.

Results in figures 2 and 3 were computed using a random perturbation about an homogeneous initial condition. However, as Turing stated, “Most of an organism, most of the time, is developing from one pattern into another, rather than from homogeneity into a pattern” [1]. To gain insights in biological development and morphogenesis, it may be useful to study Turing systems with different patterns as initial conditions. Morphogen gradients have been observed during early embryonic pattern formation in multiple studies [41, 42], suggesting a gradient in one or both morphogens might be a sensible initial pattern to consider.

A gradient was imposed along a direction with values ranging from 0 to the value (*u*_0_, *v*_0_) using the model parameters of figure 2F. Initial gradients were prescribed for either *v* (figure 4A, top row), *u* (figure 4B, top row) or for both (figure 4C, top row). The initial conditions on *u* have a stronger impact than initial conditions on *v*. When an initial gradient was imposed for *v*, the final pattern was similar to the pattern obtained for random initial conditions, which corresponds to mode (1, 1, 1) (figure 4A, bottom row). However, an initial gradient on *u*, regardless of the initial values considered for *v*, resulted in a final pattern associated with mode (0, 0, 2) (figure 4B and C, bottom row). This stronger effect of the reactant *u* may be explained by its higher exponent in the term *u*^2^*v* with respect to *v* in the nonlinear reaction term (see equation (2)). Imposing a specific pattern as initial condition allows in this case targeting a particular mode.

**Figure 4:**
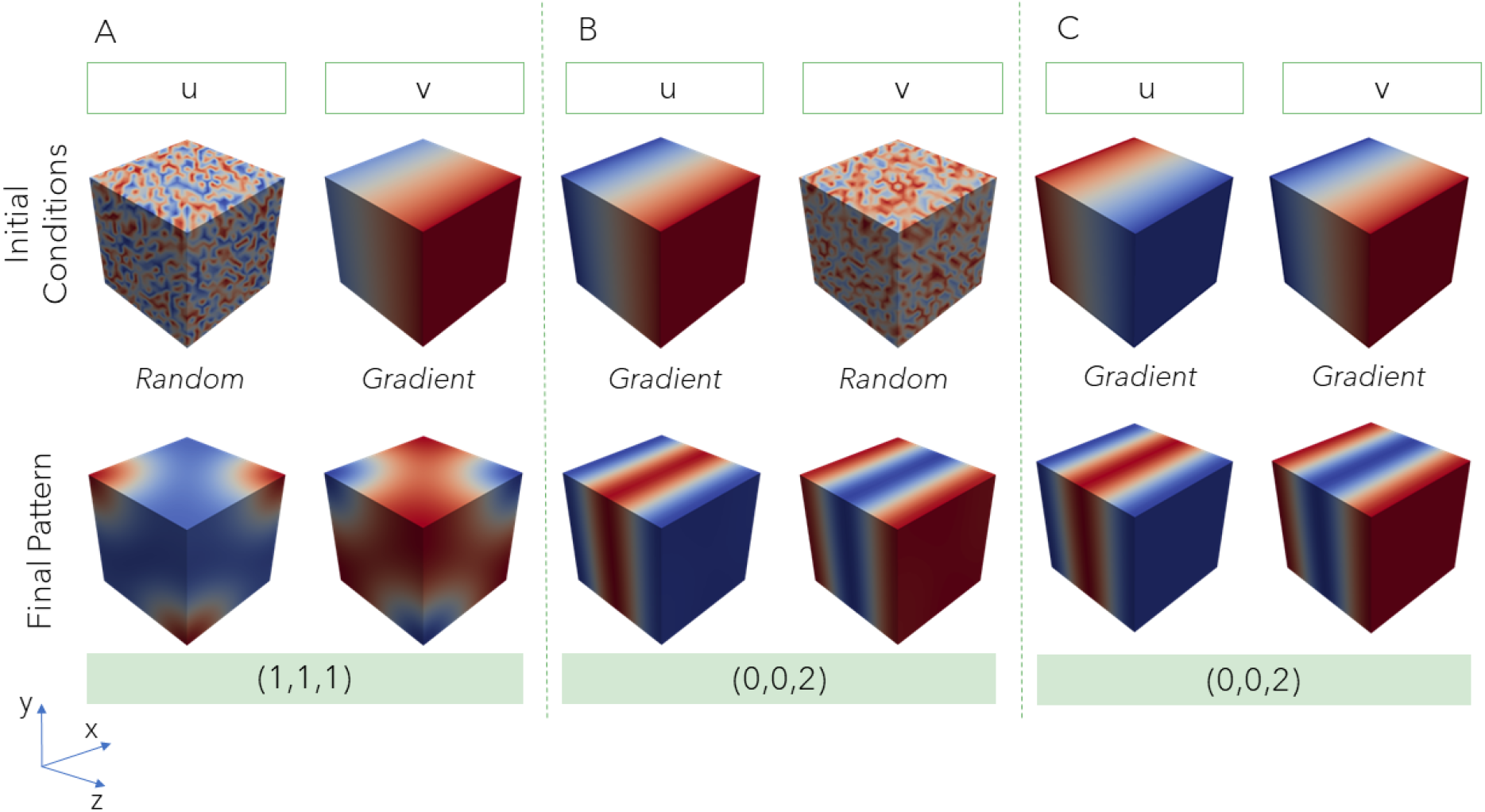
Effect of imposing gradients as initial conditions on the final pattern. The same parameters and dimensions as in figure 2F are used. Red represents the highest values of the reactant concentration while blue corresponds to the lowest ones. The initial conditions considered are shown in the top row while final patterns are shown in the bottom row. Note that Schnakenberg kinetics result in the two reactants being out of phase. An initial gradient for *v* and random values for *u* (A) result in the same final pattern (corresponding to mode (1, 1, 1)) as having both *u* and *v* with random initial values (figure 2F). However, an initial gradient for *u* results in a final pattern corresponding to mode (0, 0, 2) for both initial random values (B) and an initial gradient (C) in *v*.

### 3.4 Influence of a growing domain on the final patterns

Results in the previous sections reveal the influence of domain size on the admissible modes, in addition to the significant role of initial conditions in mode competition. Thus, emerging patterns are dependant on domain dimensions and possibly existing pre-patterning. Since template size increases during most developmental events such as gastrulation or limb formation, it follows that morphogen pattern evolution will likely be determined, in part, by the growth of the domain.

We conducted numerical simulations to examine the effect of a growing domain on the resulting patterns. To model domain growth, we took the converged pattern of a first simulation in a cubic domain starting from random initial conditions (figure 2F) and imposed this pattern as initial conditions in an elongated geometry, where the mesh size was increased by 10% along a direction (represented by a dotted arrow in figure 5). Next, we ran the simulation with this new domain size and the prescribed initial conditions until the pattern converged once again (represented by a solid arrow in figure 5). This process was repeated three more times, increasing the domain size along the same direction by 10% each time, until we reached a total elongation of 40%. The top rows of figure 5 show the initial conditions considered and the converged patterns obtained for each step in this growth process. For comparison, a pattern obtained for the same domain size in each step of the process, but considering random initial conditions, is shown in the bottom rows of figure 5. Note the difference in converged patterns obtained at each step of the growth process with respect to the equivalent domain size with random initial conditions.

**Figure 5:**
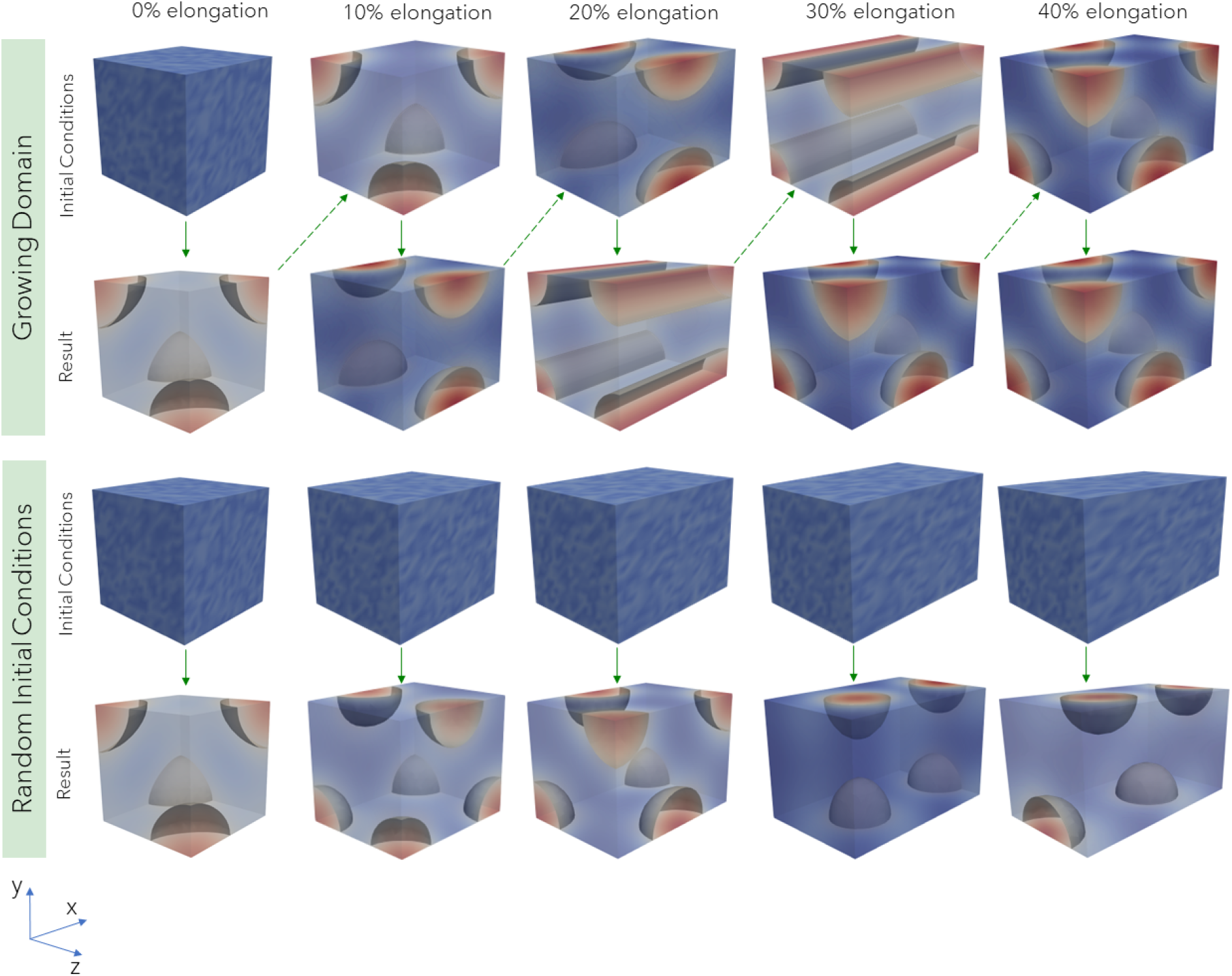
Effect of a growing domain on the final pattern of u. Red represents the highest activator values and blue the lowest ones. The initial conditions are shown above the final patterns for all simulations. For the growing domain, initial conditions correspond to the final pattern of the previous step, stretched by 10% along a direction (top two rows). In the other cases, each simulation is computed considering a random perturbation of up to 10% about the linearised steady-state solution as initial conditions at each corresponding domain size. The vertical arrows represent the simulation process and the dotted arrows represent the stretching of the domain. The patterns obtained at each step of a growing domain are different from the ones obtained for the same domain size but starting from random initial conditions.

## 4 Discussion

The few groups that have explored Turing patterns in 3D domains to date [19, 20, 26–32] have focused on predicting and classifying the complex patterns arising due to the added dimension. In contrast, our study shows that the pattern predicted by a Turing system is an addition of pure patterns associated with the admissible modes computed from the linearised equations. We have demonstrated that the contribution of these modes to the final pattern depends on their associated eigenvalues and the initial conditions considered. We have also explored the effect of a growing domain on pattern emergence.

### 4.1 The largest eigenvalue does not always determine the final pattern

Turing patterns can help us understand how and why certain patterns emerge in Nature. Linear stability analysis provides a tool to predict the emerging patterns through the study of the admissible modes and how they interact together. The power of this technique resides on its simplicity in both formulation and computation. However, as we have seen in this and past studies [13,18,19,25], it is limited due to the nonlinearities of the reaction-diffusion system, which can play a strong role during mode competition.

Mode contribution is determined, in part, by the eigenvalue corresponding to each admissible mode. It is often assumed that the largest eigenvalue will have the largest growth factor, and thus, will determine the dominant pattern for small random initial conditions [3]. We have shown that, even in this case, associating the largest eigenvalue and the dominant mode in the final pattern is not straightforward, especially when the eigenvalues corresponding to competing modes are close together (figure 2).

To unequivocally predict the final pattern using only linear stability analysis, we would have to select the model parameters to target only one admissible mode, which will result in what we call a pure mode, e.g., the example in figure 2E. The linear combination of patterns, as assumed in the linear analysis, is altered by nonlinear effects. Indeed, two steady solutions (*u*_1_, *v*_1_) and (*u*_2_, *v*_2_) of the Schnakenberg system in (1)-(2), if added, would yield a transient solution where

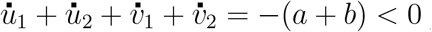

so that the system would necessary evolve and reach a different steady-state solution. Nonetheless, linear analysis has allowed us to sufficiently predict and explain most of the results simulated here numerically.

Past studies in 3D domains have shown intricate patterns arising for random initial conditions [20, 27, 29–32]. Complex patterns result from the combination of pure modes as illustrated in SI, figure S2. When several admissible modes coexist, the dominant modes cannot always be consistently predicted. We believe the reliability of the emerging pattern is associated to the proximity between the eigenvalues of the different admissible modes. When the largest eigenvalue was far from the other eigenvalues, only one pattern corresponding to the mode of the largest eigenvalue was observed for all simulations (figure 3A). However, we obtained different patterns as we repeated simulations when the eigenvalues were close together (figure 3B-F). In the latter case, the different randomness in each simulation was enough to also generate translations of a same pattern (figure 3D-F) as well as produce a pattern in which only two of the three modes sharing a same eigenvalue contributed (figure 3E).

The coexistence of multiple patterns for a same set of parameters and general conditions has been one of the main criticisms of the Turing system as a model for pattern emergence in morphogenesis [6, 43]. However, we have shown that with suitable set of parameters, leading to one eigenvalue far from the others, the final pattern is consistent and predictable when considering different random initial conditions (figure 3A).

### 4.2 Initial conditions on *u* have a stronger impact on pattern prediction than initial conditions on *v*

Initial conditions are known to play a crucial role in mode competition and, therefore, the final pattern [3, 36]. When initial conditions are similar to one of the admissible modes, this mode typically dominates in the final pattern. This is the case in figure 4C, in which the final pattern associated to mode (0, 0, 2) is the closest pattern to the gradient imposed as initial conditions in the simulation. Interestingly, due to the non-symmetric roles of activator and inhibitor in the reaction terms, initial conditions have different effects on each reactant: when an initial gradient is imposed only on the activator *u* (figure 4B) the same result is observed, while equivalent initial conditions only on *v* (figure 4A) do not seem to affect the final pattern. Turing patterns arise when a local self-enhancing reaction is coupled to a longer range antagonistic process. This may be accomplished in several ways, as described by Meinhardt [6], but always requires *v* to diffuse faster than *u*.

For the Schnakenberg kinetics (2) used in the present study, the activator *u* fulfils the role of self-activation through a nonlinear positive feedback on itself, but also inhibits the production of *v*. In turn, *v* is self-inhibiting but with a linear feedback on itself and acts as an activator of *u*. Then, the concentration of *v* decreases in response to high concentrations of *u*. The quadratic term *u*^2^ amplifies the effect of the activator, likely making it more sensitive to changes in its concentration. In contrast, the effect of *v* is linear, which means that its influence is proportional to its concentration. Altogether, it seems reasonable that initial conditions on *u* have a strong influence on the final pattern while initial conditions on *v* do not.

A similar dependence on the initial concentration of *u* has been reported for the Gierer-Meinhardt [39] model. A small baseline production of activator *u* initiates patterning even at low concentrations, whereas a small baseline production of inhibitor *v* can maintain a stable inactive state until activated by an external trigger like an influx of activator from a neighbouring zone [6]. Hence, it is plausible to assume that similar behaviours will be observed for all models in which the reactant *u* promotes pattern emergence. Nonetheless, whether imposing specific initial conditions on either reactant for different kinetic models affects the resulting pattern merits further study. Future work will also include testing alternative patterns as initial conditions, such as a local source of reactant, to verify whether such asymmetry in pattern emergence is still observed.

### 4.3 Bifurcations and path dependence in a growing domain

The Turing system provides a useful model for understanding how patterns arise during a variety of morphogenetic processes, which generally involve a growing structure. Morphogen expression driving these processes evolves in the expanding tissue over time. Hence, studying the emergence of Turing patterns in 3D growing domains may provide insights into the factors and conditions involved in morphogenesis.

As discussed in the previous section, initial conditions can enhance certain admissible modes when they are close to the pattern associated with one of these modes. However, the admissible modes may change within a growing domain, such that some modes might appear and others disappear as the domain increases in size, giving rise to bifurcation events in the configuration space. This may condition how the observed pattern evolves, which could explain the predictions at different stages of the growing domain in figure 5. The admissible modes associated to each domain size in this example are given in figure 6.

**Figure 6:**
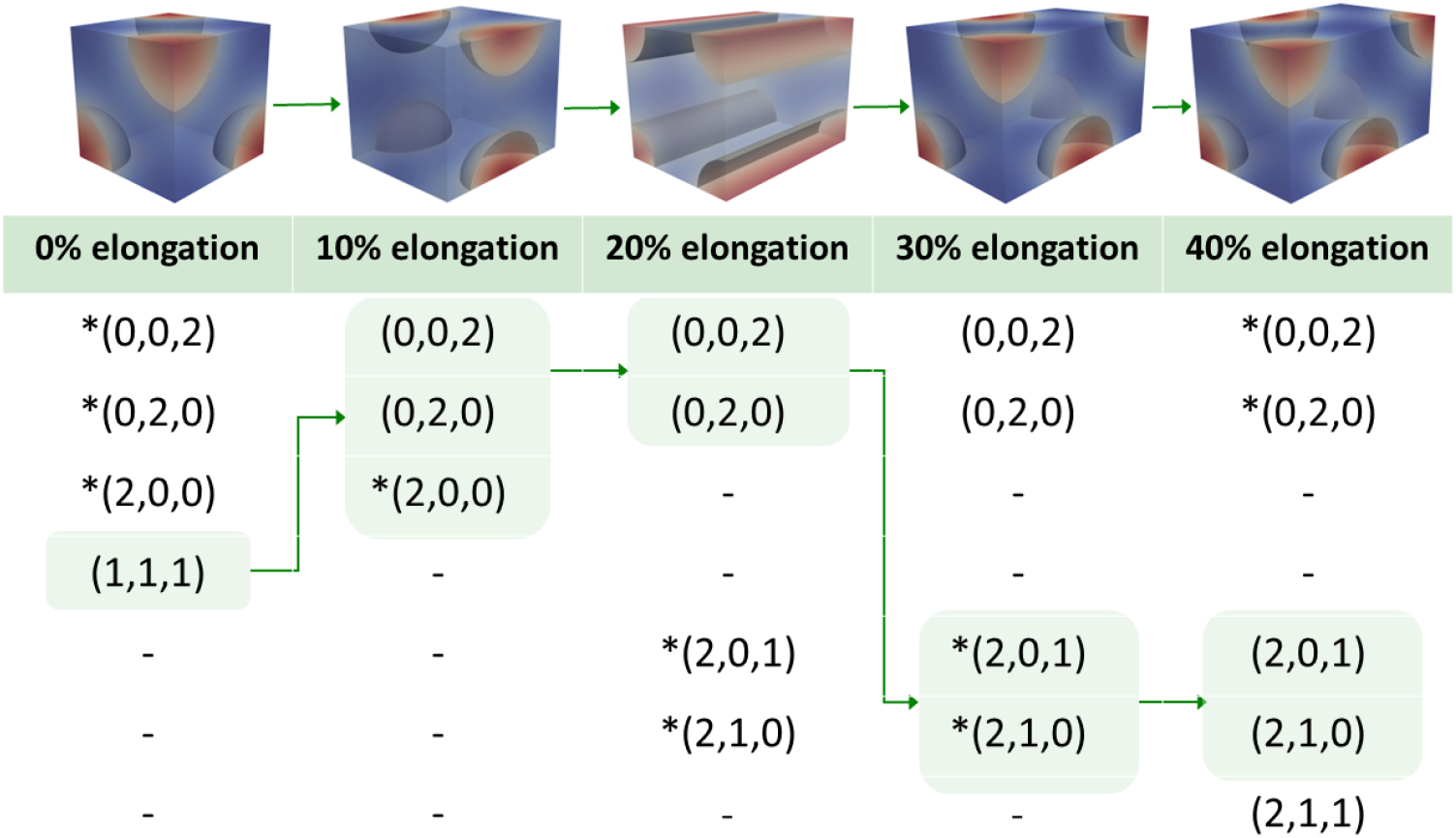
Admissible modes change as the domain grows, forcing the pattern evolution as seen in figure 5. The admissible modes identified for each domain size are listed in each column. Asterisks (*) indicate the modes with the largest eigenvalues. The header indicates the percentage of elongation of the domain. The modes associated with the pattern observed at each domain size increment have been highlighted, showing the path the pattern followed during its evolution with the growing domain.

We observe that for the domain size corresponding to 20% elongation, the final pattern for the growing domain corresponds to a combination of modes (0, 2, 0) and (2, 0, 0). Here, initial conditions were the pattern associated with the sum of modes (0, 0, 2), (0, 2, 0) and (2, 0, 0), but only the former two are admissible modes for this domain size. It seems that the resulting pattern “keeps” the admissible modes of the initial conditions, namely, (0, 0, 2)+(0, 2, 0), instead of “switching” to the new modes which have the largest eigenvalue. However, when starting from random initial conditions (figure 5), the new admissible modes (0, 1, 2) and (1, 0, 2) may be more likely to be favoured. Our simulations (SI, figure S3) show that different combinations of the admissible modes form various final patterns when starting from random initial conditions. Similar analyses can be made for the other steps in the growing domain.

A recent study already showed that domain growth in curved surfaces affects pattern selection [44]. A previous study in 1D had discussed the history dependence of pattern evolution, and the transiency of certain modes in growing domains [45]. Here, we have made similar observations in 3D and associated pattern evolution to the corresponding (transient) admissible modes. Although we were not exhaustive in the study of pattern repeatability for each domain size (SI, figure S3), and we are neglecting convective terms and growth rate effects [46], our example illustrates how changes in admissible modes may drive pattern evolution as the domain grows. In fact, our results appear to be consistent with the recent conjecture by Van Gorder et al. [46], stating that how long a given mode remains admissible within a growing domain is more determinant in pattern selection than the magnitude of its associated eigenvalue. Given that growth has been suggested as a mechanism for endowing robustness to emerging Turing patterns [34, 35], further investigations on the influence of growth rate would be of interest. Especially, because robustness of the solution has been seen to break down when growth is too fast or too slow [46]. Overall, exploring the impact of these factors as well as the effect of growth type (e.g. stretching the domain versus adding domain in an apical manner) on the reliability of final pattern prediction is especially relevant in the study of morphogenetic processes.

### 4.4 Challenges and opportunities for 3D Turing models in morphogenesis

We have begun to explore the effect of initial conditions and growth on simple 3D Turing patterns with the ultimate goal of modelling more complex patterns of morphogen expression observed in development. We explored a gradient as initial conditions because it is a morphogenic pattern frequently identified in early embryogenesis and they have been associated with the polar-axial organisation of cells [6]. For example, a gradient of retinoic acid signalling appears to coordinate the proximo-distal patterning in vertebrate limb formation [47]. Following positional information theory [48], such a concentration gradient is believed to inform cells of their relative position in space, and these then differentiate accordingly. Yet, in the past years, it has become apparent that Turing-like self-organising mechanisms may work in conjunction with cell-fate informing gradient establishment [43].

Incorporating additional components into the reaction-diffusion model has allowed investigation of biological mechanisms driving digit patterning [49, 50], embryonic axis specification [50] and frog embryo gastrulation [51], among others. Coupling reaction-diffusion systems with tissue growth [52, 53] provides additional means to explore the factors regulating these processes. The role of physical interactions between cells and how they respond to their mechanical environment have also been considered in mechanochemical models of pattern formation [10, 12]. All these mechanisms work together to generate the complex spatial patterns that define the tissues and organs of the developing organism and they are now starting to be integrated together into a more comprehensive understanding of the processes involved.

One of the main concerns in using Turing-based systems to model morphogenesis is their robustness or lack thereof. As mentioned in the previous section, domain growth under specific conditions may make pattern emergence more robust. The use of different boundary conditions for each reactant has also been shown to reduce the sensitivity of patterns to domain changes [54]. We considered homogeneous Neumann BCs, i.e. no flux conditions at the domain boundary, because we believe that, in general, it adequately represents biological conditions. However, the use of Dirichlet or mixed boundary conditions could be a closer approximation in specific morphogenetic events, which is worth exploring. Remarkably, boundary conditions do affect the number of potential symmetries in the resulting solutions, and the probability of the emerging patterns. Only through a better understanding of the fundamental elements driving the patterning in Turing models, can we successfully use them to elucidate the biophysical mechanisms in morphogenesis.

Finally, extension from 1D and 2D patterns to 3D is not trivial, and we must study 3D domains to fully capture the characteristics of certain developmental processes such as limb structure asymmetries. Predicting the type of pattern that will arise in these domains, and relating it to the model parameters and initial conditions as we have done in this study is an additional step towards developing useful models to interrogate biological hypotheses. All in all, the use of Turinglike reaction-diffusion systems to model the spatial-temporal expression of morphogens in 3D will enable us to probe the factors and conditions that affect the emergence of patterns at the organ level, which then drive cell fate, throughout the complete morphogenetic process.

## 5 Conclusions

The use of Turing patterns as a means to explore morphogen expression is widespread in developmental biology applications. However, a full understanding of the factors that lead to a specific pattern forming in three-dimensional (3D) domains is still lacking. Through linear stability analysis and finite element modelling, we have associated the admissible modes of the linearised Turing system with the emerging patterns, and studied the effects of initial conditions and domain growth in 3D.

Our results reveal that nonlinearities play a strong role when the eigenvalues of the linearised system are close to each other. This can lead to less robust predictions, with different patterns emerging for repeated simulations in which random perturbations around the linearised steadystate solution are considered as initial conditions. We also demonstrate that the effect of initial conditions is asymmetric between the reactants *u* and *v*, with initial conditions on *u* having a greater influence than the initial conditions on *v*. Therefore, carefully selecting the model parameters that determine the system’s eigenvalues and the initial conditions are crucial factors in accurately predicting 3D Turing patterns. Finally, the bifurcations in admissible modes that we have seen within a growing domain have the potential to improve the reliability of predictions, which supports the use of Turing patterns as a model for morphogenesis.

Further research on Schnakenberg and other alternative kinetic models is necessary to gain a complete understanding of how 3D Turing patterns evolve in growing domains, as this particular field of research has been relatively overlooked. Exploring the emergence of patterns, and their dependence on the evolution of domain size is particularly relevant to the study of developmental processes such as limb formation. By combining computational modelling of Turing patterns with experimental analyses of morphogen expression, we will improve our understanding of morphogen patterning and gain further insights into the mechanisms that underlie these processes.

## Data Accessibility

The code used to generate the finite element results in this study can be accessed at https://gitlab.com/turing-embryogenesis-group/TuringPattern. All other data required for reproducing the results in this study are included in the article and/or supporting information.

## Acknowledgements

This work was completed using the Discovery cluster, supported by Northeastern University’s Research Computing team.

## References

[1] Turing AM. The chemical basis of morphogenesis. Philosophical Transactions of the Royal Society of London. 1952;237(641):37–72. Available from: https://doi.org/10.1007/BF02459572. doi: 10.1007/BF02459572.

[2] Harrison LG. Kinetic theory of living pattern. Developmental and Cell Biology Series; 28. Cambridge: Cambridge University Press; 1993. Available from: https://doi.org/10.1017/CBO9780511529726. doi: 10.1017/CBO9780511529726.

[3] Murray JD. Mathematical Biology II: Spatial Models and Biomedical Applications. vol. 18 of Interdisciplinary Applied Mathematics. 3rd ed. New York, NY: Springer New York; 2003. Available from: http://link.springer.com/10.1007/b98869. doi: 10.1007/b98869.

[4] Meinhardt H, Gierer A. Pattern formation by local self-activation and lateral inhibition. BioEssays. 2000;22(8):753–760. Available from: https://doi.org/10.1002/1521-1878(200008)22:8%3C753::AID-BIES9%3E3.0.CO;2-Z. doi: 10.1002/1521-1878(200008)22:8%753::AID-BIES9%3.0.CO;2-Z.

[5] Miura T, Shiota K, Morriss-Kay G, Maini PK. Mixed-mode pattern in Doublefoot mutant mouse limb – Turing reaction–diffusion model on a growing domain during limb development. Journal of Theoretical Biology. 2006;240(4):562–573. Available from: https://doi.org/10.1016/j.jtbi.2005.10.016. doi: 10.1016/j.jtbi.2005.10.016.

[6] Meinhardt H. Turing’s theory of morphogenesis of 1952 and the subsequent discovery of the crucial role of local self-enhancement and long-range inhibition. Interface Focus. 2012;2(4):407–416. Available from: https://doi.org/10.1098/rsfs.2011.0097. doi: 10.1098/rsfs.2011.0097.

[7] Marcon L, Sharpe J. Turing patterns in development: What about the horse part? Current Opinion in Genetics and Development. 2012;22(6):578–584. Available from: https://doi.org/10.1016/j.gde.2012.11.013. doi: 10.1016/j.gde.2012.11.013.

[8] Economou AD, Ohazama A, Porntaveetus T, Sharpe PT, Kondo S, Basson MA, et al. Periodic stripe formation by a Turing mechanism operating at growth zones in the mammalian palate. Nature Genetics. 2012;44(3):348–351. Available from: https://doi.org/10.1038/ng.1090. doi: 10.1038/ng.1090.

[9] Menshykau D, Michos O, Lang C, Conrad L, McMahon AP, Iber D. Image-based modeling of kidney branching morphogenesis reveals GDNF-RET based Turing-type mechanism and pattern-modulating WNT11 feedback. Nature Communications. 2019;10(1):239. Available from: https://doi.org/10.1038/s41467-018-08212-8. doi: 10.1038/s41467-018-08212-8.

[10] Rueda-Contreras MD, Romero-Arias JR, Aragon JL, Barrio RA. Curvature-driven spatial patterns in growing 3D domains: A mechanochemical model for phyllotaxis. PloS ONE. 2018;13(8):e0201746. Available from: https://doi.org/10.1371/journal.pone.0201746. doi: 10.1371/journal.pone.0201746.

[11] Okuda S, Miura T, Inoue Y, Adachi T, Eiraku M. Combining Turing and 3D vertex models reproduces autonomous multicellular morphogenesis with undulation, tubulation, and branching. Scientific Reports. 2018;8(1):1–15. Available from: https://doi.org/10.1038/s41598-018-20678-6. doi: 10.1038/s41598-018-20678-6.

[12] Brinkmann F, Mercker M, Richter T, Marciniak-Czochra A. Post-Turing tissue pattern formation: Advent of mechanochemistry. PLoS Computational Biology. 2018;14(7):1–21. Available from: https://doi.org/10.1371/journal.pcbi.1006259. doi: 10.1371/journal.pcbi.1006259.

[13] Borckmans P, De Wit A, Dewel G. Competition in ramped Turing structures. Physica A: Statistical Mechanics and its Applications. 1992;188(1-3):137–157. Available from: https://doi.org/10.1016/0378-4371(92)90261-N. doi: 10.1016/0378-4371(92)90261-N.

[14] Lyons MJ, Harrison LG. Stripe selection: An intrinsic property of some pattern-forming models with nonlinear dynamics. Developmental Dynamics. 1992;195(3):201–215. Available from: https://doi.org/10.1002/aja.1001950306. doi: 10.1002/aja.1001950306.

[15] Barrio RA, Varea C, Aragón JL, Maini PK. A two-dimensional numerical study of spatial pattern formation in interacting Turing systems. Bulletin of Mathematical Biology. 1999;61(3):483–505. Available from: https://doi.org/10.1006/bulm.1998.0093. doi: 10.1006/bulm.1998.0093.

[16] Yang L, Dolnik M, Zhabotinsky AM, Epstein IR. Spatial Resonances and Superposition Patterns in a Reaction-Diffusion Model with Interacting Turing Modes. Physical Review Letters. 2002;88(20):4. Available from: https://doi.org/10.1103/PhysRevLett.88.208303. doi: 10.1002/aja.1001950306.

[17] Barrio RA, Maini PK, Aragón JL, Torres M. Size-dependent symmetry breaking in models for morphogen-esis. Physica D: Nonlinear Phenomena. 2002;168-169:61–72. Available from: https://doi.org/10.1016/S0167-2789(02)00495-5. doi: 10.1016/S0167-2789(02)00495-5.

[18] Ermentrout B. Stripes or spots? Nonlinear effects in bifurcation of reaction-diffusion equations on the square. Proceedings of the Royal Society of London Series A: Mathematical and Physical Sciences. 1991;434(1891):413–417. Available from: https://doi.org/10.1098/rspa.1991.0100. doi: 10.1098/rspa.1991.0100.

[19] Leppänen T, Karttunen M, Barrio RA, Kaski K. Morphological transitions and bistability in Turing systems. Physical Review E. 2004;70(6):9. Available from: https://doi.org/10.1103/PhysRevE.70.066202. doi: 10.1103/PhysRevE.70.066202.

[20] Shoji H, Yamada K. Most stable patterns among three-dimensional Turing patterns. Japan Journal of Industrial and Applied Mathematics. 2007;24:67–77. Available from: https://doi.org/10.1007/BF03167508. doi: 10.1007/BF03167508.

[21] Wei J, Winter M. Stationary multiple spots for reaction-diffusion systems. Journal of Mathematical Biology. 2008;57(1):53–89. Available from: https://doi.org/10.1007/s00285-007-0146-y. doi: 10.1007/s00285-007-0146-y.

[22] Liu P, Shi J, Wang Y, Feng X. Bifurcation analysis of reaction-diffusion Schnakenberg model. Journal of Mathematical Chemistry. 2013;51(8):2001–2019. Available from: https://doi.org/10.1007/s10910-013-0196-x. doi: 10.1007/s10910-013-0196-x.

[23] Chen Y, Buceta J. A non-linear analysis of Turing pattern formation. PLoS ONE. 2019;14(8):e0220994. Available from: https://doi.org/10.1371/journal.pone.0220994. doi: 10.1371/journal.pone.0220994.

[24] Krause AL, Klika V, Woolley TE, Gaffney EA. From one pattern into another: Analysis of Turing patterns in heterogeneous domains via WKBJ. Journal of the Royal Society Interface. 2020;17(162):20190621. Available from: https://doi.org/10.1098/rsif.2019.0621. doi: 10.1098/rsif.2019.0621.

[25] Khudhair HK, Zhang Y, Fukawa N. Pattern selection in the Schnakenberg equations: From normal to anomalous diffusion. Numerical Methods for Partial Differential Equations. 2021;38:1843–1860. Available from: https://doi.org/10.1002/num.22842. doi: 10.1002/num.22842.

[26] De Wit A, Dewel G, Borckmans P, Walgraef D. Three-dimensional dissipative structures in reaction-diffusion systems. Physica D: Nonlinear Phenomena. 1992;61(1-4):289–296. Available from: https://doi.org/10.1016/0167-2789(92)90173-K. doi: 10.1016/0167-2789(92)90173-K.

[27] De Wit A, Borckmans P, Dewel G. Twist grain boundaries in three-dimensional lamellar Turing structures. Proceedings of the National Academy of Sciences of the United States of America. 1997;94(24):12765–12768. Available from: https://doi.org/10.1073/pnas.94.24.12765. doi: 10.1073/pnas.94.24.12765.

[28] Callahan TK, Knobloch E. Pattern formation in three-dimensional reaction-diffusion systems. Physica D. 1999;132(3):339–362. Available from: https://doi.org/10.1016/S0167-2789(99)00041-X. doi: 10.1016/S0167-2789(99)00041-X.

[29] Leppänen T, Karttunen M, Kaski K, Barrio RA, Zhang L. A new dimension to Turing patterns. Physica D: Nonlinear Phenomena. 2002;168-169:35–44. Available from: https://doi.org/10.1016/S0167-2789(02)00493-1. doi: 10.1016/S0167-2789(02)00493-1.

[30] Leppänen T, Karttunen M, Kaski K, Barrio RA. Dimensionality effects in Turing pattern formation. International Journal of Modern Physics B. 2003;17(29):5541–5553. Available from: https://doi.org/10.1142/S0217979203023240. doi: 10.1142/S0217979203023240.

[31] Shoji H, Yamada K, Ohta T. Interconnected Turing patterns in three dimensions. Physical Review E. 2005;72(6):1–4. Available from: https://doi.org/10.1103/PhysRevE.72.065202. doi: 10.1103/Phys-RevE.72.065202.

[32] Shoji H, Ohta T. Computer simulations of three-dimensional Turing patterns in the Lengyel-Epstein model. Physical Review E. 2015;91(3):1–11. Available from: https://doi.org/10.1103/PhysRevE.91.032913. doi: 10.1103/PhysRevE.91.032913.

[33] Song W, Wubs F, Thies J, Baars S. Numerical bifurcation analysis of a 3D turing-type reaction–diffusion model. Communications in Nonlinear Science and Numerical Simulation. 2018;60:145–164. Available from: https://doi.org/10.1016/j.cnsns.2018.01.003. doi: 10.1016/j.cnsns.2018.01.003.

[34] Barrass I, Crampin EJ, Maini PK. Mode transitions in a model reaction–diffusion system driven by domain growth and noise. Bulletin of Mathematical Biology. 2006;68:981–995. Available from: https://doi.org/10.1007/s11538-006-9106-8. doi: 10.1007/s11538-006-9106-8.

[35] Maini PK, Woolley TE, Baker RE, Gaffney EA, Seirin Lee S. Turing’s model for biological pattern formation and the robustness problem. Interface Focus. 2012 aug;2(4):487–496. Available from: https://doi.org/10.1098/rsfs.2011.0113. doi: 10.1098/rsfs.2011.0113.

[36] Hernandez-Aristizabal D, Garzón-Alvarado DA, Madzvamuse A. Turing Pattern Formation Under Heterogen-eous Distributions of Parameters for an Activator-Depleted Reaction Model. Journal of Nonlinear Science. 2021;31(2). Available from: https://doi.org/10.1007/s00332-021-09685-6. doi: 10.1007/s00332-021-09685-6.

[37] Schnakenberg J. Simple chemical reaction systems with limit cycle behaviour. Journal of Theoretical Biology. 1979;81(3):389–400. Available from: https://doi.org/10.1016/0022-5193(79)90042-0. doi: 10.1016/0022-5193(79)90042-0.

[38] Zhu M, Murray JD. Parameter domains for generating spatial pattern: comparison of reaction-diffusion and cell-chemotaxis models. International Journal of Bifurcation and Chaos. 1995;5(6):1503–1524. Available from: https://doi.org/10.1142/S0218127495001150. doi: 10.1142/S0218127495001150.

[39] Gierer A, Meinhardt H. A theory of biological pattern formation. Kybernetik. 1972;12(1):30–39. Available from: https://doi.org/10.1007/BF00289234. doi: 10.1007/BF00289234.

[40] Iron D, Wei J, Winter M. Stability analysis of Turing patterns generated by the Schnakenberg model. Journal of Mathematical Biology. 2004;49(4):358–390. Available from: http://doi.org/10.1007/s00285-003-0258-y. doi: 10.1007/s00285-003-0258-y.

[41] Nüsslein-Volhard C, Wieschaus E. Mutations affecting segment number and polarity in Drosophila. Nature. 1980;287(5785):795–801. Available from: https://doi.org/10.1038/287795a0. doi: 10.1038/287795a0.

[42] Balasubramanian R, Zhang X. Mechanisms of FGF gradient formation during embryogenesis. Seminars in Cell & Developmental Biology. 2016;53:94–100. Available from: https://doi.org/10.1016/j.semcdb.2015.10.004. doi: 10.1016/j.semcdb.2015.10.004.

[43] Green JBA, Sharpe J. Positional information and reaction-diffusion: Two big ideas in developmental biology combine. Development. 2015;142(7):1203–1211. Available from: https://doi.org/10.1242/dev.114991. doi: 10.1242/dev.114991.

[44] Sánchez-Garduño F, Krause AL, Castillo JA, Padilla P. Turing–Hopf patterns on growing domains: The torus and the sphere. Journal of Theoretical Biology. 2019 nov;481:136–150. Available from: https://doi.org/10.1016/j.jtbi.2018.09.028. doi: 10.1016/j.jtbi.2018.09.028.

[45] Klika V, Gaffney EA. History dependence and the continuum approximation breakdown: The impact of domain growth on Turing’s instability. Proceedings of the Royal Society A: Mathematical, Physical and Engineering Sciences. 2017;473(2199):20160744. Available from: https://doi.org/10.1098/rspa.2016.0744. doi: 10.1098/rspa.2016.0744.

[46] Van Gorder RA, Klika V, Krause AL. Turing conditions for pattern forming systems on evolving manifolds. Journal of Mathematical Biology. 2021;82(1-2):4. Available from: https://doi.org/10.1007/s00285-021-01552-y. doi: 10.1007/s00285-021-01552-y.

[47] McQueen C, Towers M. Establishing the pattern of the vertebrate limb. Development. 2020;147(17). Available from: https://doi.org/10.1242/dev.177956. doi: 10.1242/dev.177956.

[48] Wolpert L. Positional information revisited. Development. 1989;107(SUPPL.):3–12. Available from: https://doi.org/10.1242/dev.107.supplement.3. doi: 10.1242/dev.107.supplement.3.

[49] Raspopovic J, Marcon L, Russo L, Sharpe J. Digit patterning is controlled by a Bmp-Sox9-Wnt Turing network modulated by morphogen gradients. Science. 2014;345(6196):566–570. Available from: https://doi.org/10.1126/science.1252960. doi: 10.1126/science.1252960.

[50] Marcon L, Diego X, Sharpe J, Müller P. High-throughput mathematical analysis identifies Turing networks for patterning with equally diffusing signals. eLife. 2016;5:e14022. Available from: https://doi.org/10.7554/eLife.14022. doi: 10.7554/eLife.14022.

[51] Nesterenko AM, Kuznetsov MB, Korotkova DD, Zaraisky AG. Morphogene adsorption as a Turing instability regulator: Theoretical analysis and possible applications in multicellular embryonic systems. PLoS ONE. 2017;12(2):1–22. Available from: https://doi.org/10.1371/journal.pone.0171212. doi: 10.1371/journal.pone.0171212.

[52] Márquez-Flórez KM, Monaghan JR, Shefelbine SJ, Ramirez-Martinez A, Garzón-Alvarado DA. A computational model for the joint onset and development. Journal of Theoretical Biology. 2018;454:345–356. Available from: https://doi.org/10.1016/j.jtbi.2018.04.015. doi: 10.1016/j.jtbi.2018.04.015.

[53] Seirin Lee S, Gaffney EA, Baker RE. The Dynamics of Turing Patterns for Morphogen-Regulated Growing Domains with Cellular Response Delays. Bulletin of Mathematical Biology. 2011;73(11):2527–2551. Available from: https://doi.org/10.1007/s11538-011-9634-8. doi: 10.1007/s11538-011-9634-8.

[54] Dillon R, Maini PK, Othmer HG. Pattern formation in generalized Turing systems: I. Steady-state patterns in systems with mixed boundary conditions. Journal of Mathematical Biology. 1994;32(4):345–393. Available from: https://doi.org/10.1007/BF00160165. doi: 10.1007/BF00160165.4

