## Supplementary material for "Turing pattern prediction in three-dimensional domains: the role of initial conditions and growth": SUPP

#### Contents

**S1 Suppl. text: Parameter values required for Turing pattern emergence** **2**

**S2 Suppl. text: Finite Element implementation** **4**

**S3 Suppl. text: Patterns obtained by adding pure modes** **6**

**S4 Suppl. figure: Complex patterns arise from a combination of modes** **7**

**S5 Suppl. figures: Admissible modes change in a growing domain** **8**

### S1 Suppl. text: Parameter values required for Turing pattern emergence

Whether the reaction-diffusion system with Schnakenberg [1] kinetics used in this study is capable of generating Turing-type spatial patterns [2] depends on the value of the dimensionless parameters $a$, $b$ and $d$. Recall that the system is given by

$$\left\{ \begin{array}{lcl} \dot{u} & = & \gamma f(u,v) + \Delta u \\ \dot{v} & = & \gamma g(u,v) + d\Delta v \end{array} \right\} \quad \text{with} \quad \left\{ \begin{array}{lcl} f(u,v) & = & a - u + u^2 v \\ g(u,v) & = & b - u^2 v \end{array} \right\}, \tag{1}$$

where $u(\boldsymbol{x},t)$ and $v(\boldsymbol{x},t)$ are the reactant concentrations. Their time derivatives are indicated by a superimposed dot and $\Delta(\bullet) = (\partial_x^2 + \partial_y^2 + \partial_z^2)(\bullet)$ is the standard Laplacian operator. The vector $\boldsymbol{x}$ indicates spatial position and $t$ is the temporal variable. The term $\gamma$ can have multiple interpretations linked to the domain size and to the relative strength of the reaction terms, and $d$ is the ratio of diffusion coefficients of the reactants. The Schnakenberg parameters $a$ and $b$ must be positive.

Our goal here is to have a homogeneous steady state which is stable in response to small perturbations in the absence of diffusion, but unstable to small spatial perturbations when diffusion is present [3]. Because the instability we are studying is diffusion-driven, the linear stability analysis is performed only for a perturbation in space. This results in the following linearised system

$$\begin{bmatrix} \dot{u} \\ \dot{v} \end{bmatrix} = \gamma \boldsymbol{A} \begin{bmatrix} u(t) \\ v(t) \end{bmatrix} + \begin{bmatrix} \Delta & 0 \\ 0 & d\Delta \end{bmatrix} \begin{bmatrix} u \\ v \end{bmatrix} \quad \text{with} \quad \boldsymbol{A} = \begin{bmatrix} f_u & f_v \\ g_u & g_v \end{bmatrix}, \tag{2}$$

where $\boldsymbol{A}$ is the stability matrix of the kinetic functions $f$ and $g$ at the steady state $(u_0, v_0)$, such that $f(u_0, v_0) = g(u_0, v_0) = 0$, and $f_u$, $f_v$, $g_u$ and $g_v$ are the derivatives evaluated at $(u_0, v_0)$. We first study the linearised system (2) without diffusion. The necessary and sufficient condition to guarantee the linear stability without diffusion is

$$\text{tr}(\boldsymbol{A}) = f_u + g_v < 0 \quad \text{and} \quad \det(\boldsymbol{A}) = f_u g_v - f_v g_u > 0. \tag{3}$$

The steady-state solution depends on the kinetic parameters $a$ and $b$. In consequence, these inequalities provide a range of parameter values for which the solutions are possible, namely,

$$b - a < (a+b)^3. \tag{4}$$

The next step is to establish the conditions to produce an instability in response to small spatial perturbations when diffusion is present. To solve the linearised system (2), we first introduce a solution of the form

$$\begin{bmatrix} u(\boldsymbol{x},t) - u_0 \\ v(\boldsymbol{x},t) - v_0 \end{bmatrix} = \boldsymbol{w}(\boldsymbol{x},t) = \sum_k c_k e^{\lambda t} \boldsymbol{w}_k(\boldsymbol{x}), \tag{5}$$

where $\boldsymbol{w}_k$ is the eigenvector corresponding to the wavevector $\boldsymbol{k}$, and $c_k$ are constants determined from a Fourier expansion of the initial conditions in terms of $\boldsymbol{w}_k(\boldsymbol{x})$. By assuming eigenvectors with the oscillatory form $\boldsymbol{w}_k(\boldsymbol{x}) = \boldsymbol{w}_0 e^{i\boldsymbol{k}\cdot\boldsymbol{x}}$, the system in (2) may be written for each wavevector as,

$$\lambda \boldsymbol{w}_k(\boldsymbol{x}) = \boldsymbol{M}\boldsymbol{w}_k(\boldsymbol{x}) \quad \text{with} \quad \boldsymbol{M} = \gamma\boldsymbol{A} - \boldsymbol{D}k^2 \;\; ; \; \boldsymbol{D} = \begin{bmatrix} 1 & 0 \\ 0 & d \end{bmatrix} \quad \text{and} \quad k = ||\boldsymbol{k}||. \tag{6}$$

Therefore, in order to ensure the system's instability, we must have that $\lambda$ in (5), i.e. the eigenvalues of $\boldsymbol{M}$, must have a positive real part. Since $tr(\boldsymbol{M}) = \gamma\, tr(\boldsymbol{A}) - k^2(1+d) < 0$, in order to have $Re(\lambda) > 0$ we need

$$\det(\boldsymbol{M}) = dk^4 - \gamma(df_u + g_v)k^2 + \gamma^2 \det(\boldsymbol{A}) \; < \; 0, \tag{7}$$

leading to the following parameter space for $d$,

$$\begin{aligned} &df_u + g_v > 0, \\ &d^2 f_u^2 + 2\left(2 f_v g_u - f_u g_v\right) d + g_v^2 > 0. \end{aligned} \tag{8}$$

In summary, to ensure the emergence of Turing patterns, the model parameters $a$, $b$ and $d$ must be selected to be in the Turing space of parameters, i.e. they must satisfy the inequalities (4) and (8). Additionally, for a given parallelepiped with dimensions $L_x \times L_u \times L_z$, the existence of integers $(m, n, p)$ that satisfy the inequalities in equation (7) of the Main Text is also necessary.

### References


[1] Schnakenberg J. Simple chemical reaction systems with limit cycle behaviour. Journal of Theoretical Biology. 1979;81(3):389–400. Available from: https://doi.org/10.1016/0022-5193(79)90042-0. doi: 10.1016/0022-5193(79)90042-0.

[2] Turing AM. The chemical basis of morphogenesis. Philosophical Transactions of the Royal Society of London. 1952;237(641):37–72. Available from: https://doi.org/10.1007/BF02459572. doi: 10.1007/BF02459572.

[3] Murray JD. Mathematical Biology II: Spatial Models and Biomedical Applications. vol. 18 of Interdisciplinary Applied Mathematics. 3rd ed. New York, NY: Springer New York; 2003. Available from: http://link.springer.com/10.1007/b98869. doi: 10.1007/b98869.

#### S2 Suppl. text: Finite Element implementation

To solve the nonlinear system of equations (1) we discretised them in space by applying the finite element method and in time using the implicit mid-point rule.

The strong form of the problem is

$$\dot{\boldsymbol{u}} = \boldsymbol{D}\Delta\boldsymbol{u} + \gamma\boldsymbol{f}(\boldsymbol{u}) \qquad \text{where} \tag{9}$$

$$\boldsymbol{u} = \begin{bmatrix} u(\boldsymbol{x},t) \\ v(\boldsymbol{x},t) \end{bmatrix}, \quad \boldsymbol{D} = \begin{bmatrix} 1 & 0 \\ 0 & d \end{bmatrix} \quad \text{and} \quad \boldsymbol{f}(\boldsymbol{u}) = \begin{bmatrix} f(u,v) \\ g(u,v) \end{bmatrix} = \begin{bmatrix} -u \\ 0 \end{bmatrix} + \begin{bmatrix} a+u^2v \\ b-u^2v \end{bmatrix}. \tag{10}$$

The kinetic functions $\boldsymbol{f}(\boldsymbol{u})$ have been split into their linear and nonlinear parts, $\boldsymbol{f}_l = [-u,\ 0]^T$ and $\boldsymbol{f}_{nl} = [a+u^2v,\ b-u^2v]^T$, respectively. Applying the *method of weighted residuals*, (9) becomes

$$\int_\Omega \dot{\boldsymbol{u}}\cdot\boldsymbol{q}\,\mathrm{d}\Omega = \int_\Omega \boldsymbol{D}\Delta\boldsymbol{u}\cdot\boldsymbol{q}\,\mathrm{d}\Omega + \int_\Omega \gamma\boldsymbol{f}_l(\boldsymbol{u})\cdot\boldsymbol{q}\,\mathrm{d}\Omega \int_\Omega \gamma\boldsymbol{f}_{nl}(\boldsymbol{u})\cdot\boldsymbol{q}\,\mathrm{d}\Omega, \forall\boldsymbol{q}\in Q \tag{11}$$

where $\boldsymbol{q}$ denotes the test functions, $\Omega$ the domain, and $Q$ a suitable functional space in $H^1$. Integrating by parts, applying homogeneous Neumann boundary conditions (e.g. the normal gradient of $\boldsymbol{u}$ is null on the contour of the domain) and reorganising the terms, the weak form in (11) reads,

$$\int_\Omega \dot{\boldsymbol{u}}\cdot\boldsymbol{q}\,\mathrm{d}\Omega + \int_\Omega \boldsymbol{D}\nabla\boldsymbol{u} : \nabla\boldsymbol{q}\,\mathrm{d}\Omega - \gamma\int_\Omega \boldsymbol{f}_l(\boldsymbol{u})\cdot\boldsymbol{q}\,\mathrm{d}\Omega - \gamma\int_\Omega \boldsymbol{f}_{nl}(\boldsymbol{u})\cdot\boldsymbol{q}\,\mathrm{d}\Omega = \boldsymbol{0}, \forall\boldsymbol{q}\in Q \tag{12}$$

is obtained. After discretisating this weak form with a set of shape functions $N_i(\boldsymbol{x})$ as $\boldsymbol{q} \approx \boldsymbol{q}^h = \sum_i N_i(\boldsymbol{x})\boldsymbol{q}_i$, and from the arbitrariness of the test functions $\boldsymbol{q}^h$, we obtain the following system of nonlinear equations

$$\mathbb{M}\boldsymbol{y} - \mathbb{K}\boldsymbol{y} + \mathbb{R}\dot{\boldsymbol{y}} - \mathbf{f_{nl}}(\boldsymbol{y}) = \boldsymbol{0} \tag{13}$$

with

$$\mathbb{M} = \begin{bmatrix} \boldsymbol{M}^b & \boldsymbol{0} \\ \boldsymbol{0} & \boldsymbol{M}^b \end{bmatrix} \text{ where } M^b_{ij} = \int_V N_i N_j \,\mathrm{d}V, \tag{14}$$

$$\mathbb{K} = \begin{bmatrix} \boldsymbol{K} & \boldsymbol{0} \\ \boldsymbol{0} & d\boldsymbol{K} \end{bmatrix} \text{ where } K_{ij} = \int_V \nabla N_i \cdot \nabla N_j \,\mathrm{d}V \tag{15}$$

$$\text{and } \mathbb{R} = \begin{bmatrix} \gamma\boldsymbol{M}^b & \boldsymbol{0} \\ \boldsymbol{0} & \boldsymbol{0} \end{bmatrix}. \tag{16}$$

The vector of unknowns $\boldsymbol{y}$ is formed by the two vectors of concentrations,

$$\boldsymbol{y} = \left[\boldsymbol{u}^h\ \boldsymbol{v}^h\right]^T, \tag{17}$$

with $\boldsymbol{u}^h = [u_1(t)\ \ u_2(t)\ldots u_n(t)]$ and $\boldsymbol{v}^h = [v_1(t)\ \ v_2(t)\ldots v_n(t)]$ the set of nodal values of the unknown fields $u$ and $v$, respectively. Also,

$$\mathbf{f}_{nl,i} = \gamma\int_V N_i \begin{bmatrix} a+u^2v \\ b-u^2v \end{bmatrix} dV \tag{18}$$

is the nonlinear reaction term at a given node $i$.

The mid-point rule is used to discretise the equations in time, resulting in

$$\mathbb{A}\,\boldsymbol{y}^{n+1} - \mathbb{B}\,\boldsymbol{y}^n - \mathbf{f}_{nl}\left(\boldsymbol{y}^{n+1/2}\right) = \boldsymbol{0}, \tag{19}$$

where $\boldsymbol{y}^{n+1/2} = \frac{1}{2}\left(\boldsymbol{y}^{n+1} + \boldsymbol{y}^{n}\right)$ and with

$$\mathbb{A} = \frac{1}{\Delta t}\mathbb{M} + \frac{1}{2}\mathbb{K} - \frac{1}{2}\mathbb{R} \quad \text{and} \quad \mathbb{B} = \frac{1}{\Delta t}\mathbb{M} - \frac{1}{2}\mathbb{K} + \frac{1}{2}\mathbb{R}. \tag{20}$$

The Newton-Raphson algorithm outlined in Algorithm 1 is used to approximate the solution to the system (19), which is nonlinear due to the term $\mathbf{f}_{nl}\left(\boldsymbol{y}^{n+1}\right)$. The matrices $\mathbb{K}$, $\mathbb{M}$ and $\mathbb{R}$ only need to be computed once before the first time step, and assembled into $\mathbb{A}$ and $\mathbb{B}$. For a constant time step size $\Delta t$, $\mathbb{A}$ and $\mathbb{B}$ will also remain unchanged at each time step.

The numerical framework outlined here has been implemented in Matlab (2022a, The MathWorks Inc.) and is available at https://gitlab.com/turing-embryogenesis-group/TuringPattern.

**Algorithm 1** Newton-Raphson algorithm to solve the nonlinear system (19) at each time step $t_n$.

1: Choose initial guess $\boldsymbol{y}_0^{n+1} = \boldsymbol{y}^n$
2: Compute residual, $\boldsymbol{r}_0 = \mathbb{A}\boldsymbol{y}_0^{n+1} - \mathbb{B}\boldsymbol{y}^n - \mathbf{f}_{nl}(\boldsymbol{y}^{n+1/2})$
3: $k = 0$
4: **while** $(\|\boldsymbol{r}\| >$ tolerance) and $(k <$ max iterations) **do**
5: $\quad k = k + 1$
6: $\quad$ Solve $(\mathbb{A} - \partial \mathbf{f}_{nl}/\partial \boldsymbol{y})\delta \boldsymbol{y} = -\boldsymbol{r}_{\boldsymbol{k+1}}$ for $\delta \boldsymbol{y}$
7: $\quad$ Update $\boldsymbol{y}_{k+1}^{n+1} = \boldsymbol{y}_k^{n+1} + \delta \boldsymbol{y}$
8: $\quad$ Compute the updated residual $\boldsymbol{r}_{k+1} = \mathbb{A}\ \boldsymbol{y}_{k+1}^{n+1} - \mathbb{B}\ \boldsymbol{y}^n - \mathbf{f}_{nl}(\boldsymbol{y}_{k+1}^{n+1/2})$
9: **end while**

### S3 Suppl. text: Patterns obtained by adding pure modes

To help interpret how pure modes predicted by the linear stability analysis combine to form the final patterns observed in the finite element simulations, we artificially generated patterns on a cubic domain.

For a parallelepipedic domain of dimensions $L_x \times L_y \times L_z$ with homogeneous Neumann boundary conditions, the solution to the linearised system for the activator $u$ at a given time $t = t^*$ can be rewritten as

$$u(\boldsymbol{x}, t^*) \simeq u_0 + \sum_{m,n,p} s_{(m,n,p)}(t^*) \cos\frac{m\pi x}{L_x} \cos\frac{n\pi y}{L_y} \cos\frac{p\pi z}{L_z} , \tag{21}$$

where $(m, n, p)$ corresponds to the *pure mode* as defined in the main text. Here, we considered only the real part of the solution and introduced the generic constant $s_{(m,n,p)}(t^*) = c_{m,n,p}\, e^{\lambda t^*}$, to simplify notation. For the case shown in figure 2A of the main text, the admissible modes are $(0, 0, 1)$, $(0, 1, 0)$ and $(1, 0, 0)$ which correspond to the solutions

$$u(\boldsymbol{x}, t^*) \simeq u_0 + s_{(0,0,1)}(t^*) \cos\frac{\pi z}{L_z} , \quad u(\boldsymbol{x}, t^*) \simeq u_0 + s_{(0,1,0)}(t^*) \cos\frac{\pi y}{L_y} ,$$
$$\text{and} \quad u(\boldsymbol{x}, t^*) \simeq u_0 + s_{(1,0,0)}(t^*) \cos\frac{\pi x}{L_x} , \tag{22}$$

respectively. Plotting these on a cubic domain produces the gradients shown in figure S1A. Then, the addition of the three modes, $(0, 0, 1)$+$(0, 1, 0)$+$(1, 0, 0)$, results in

$$u(\boldsymbol{x}, t^*) \simeq u_0 + s_{(0,0,1)}(t^*) \cos\frac{\pi z}{L_z} + s_{(0,1,0)}(t^*) \cos\frac{\pi y}{L_y} + s_{(1,0,0)}(t^*) \cos\frac{\pi x}{L_x} , \tag{23}$$

whose graphical representation for $s_{(0,0,1)}(t^*) = s_{(0,1,0)}(t^*) = s_{(1,0,0)}(t^*)$ is an eight of a sphere (figure S1B). This pattern is quite different from that of mode $(1, 1, 1)$ obtained in figure 2B of the main text. Its solution is

$$u(\boldsymbol{x}, t^*) \simeq u_0 + s_{(1,1,1)}(t^*) \cos\frac{\pi z}{L_z} \cos\frac{\pi y}{L_y} \cos\frac{\pi x}{L_x} , \tag{24}$$

which, plotted on a cubic domain, corresponds to four eights of a sphere in opposing corners of the cubic domain (figure S1C).

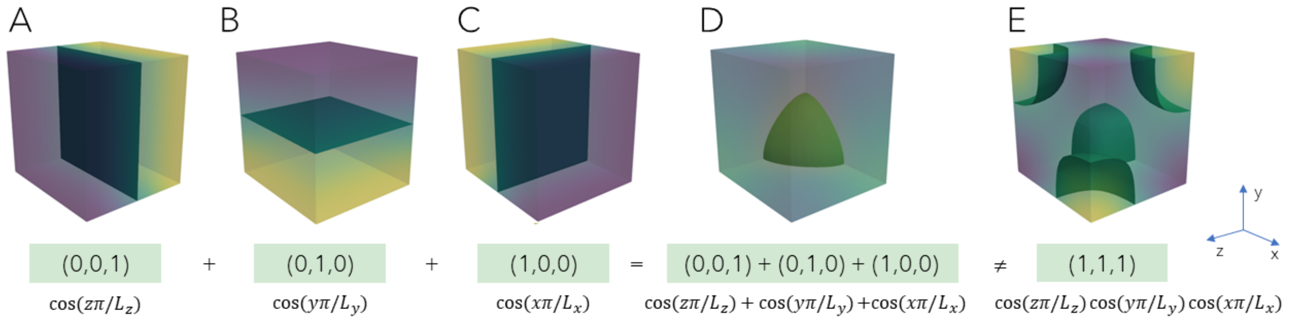


**Figure S1:** The addition of the pure modes (0,0,1), (0,1,0) and (1,0,0) results in the pattern shown in figure 2A of the main text. It is a different pattern than the pattern corresponding to mode (1,1,1), obtained in figure 2B of the main text. The patterns shown here are produced artificially for illustrative purposes.

### S4 Suppl. figure: Complex patterns arise from a combination of modes

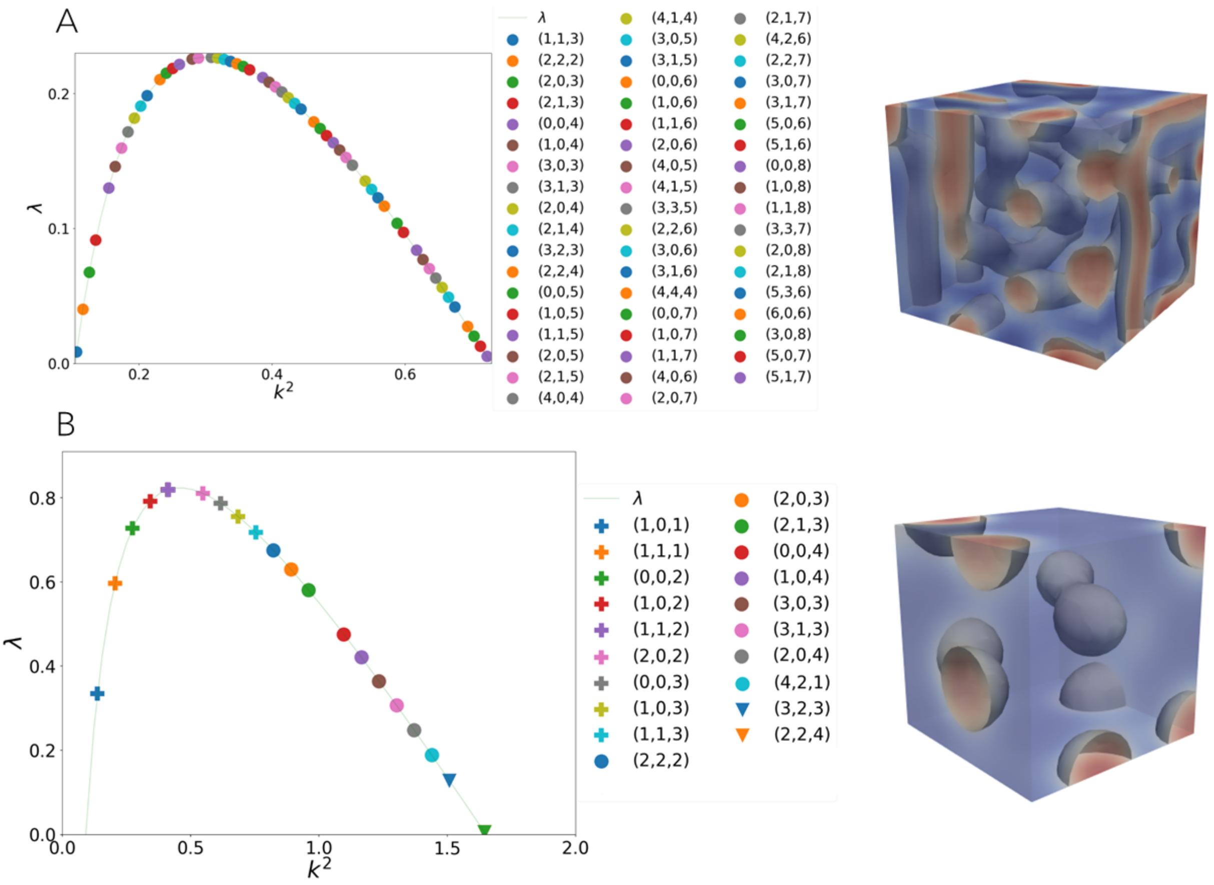


**Figure S2:** A complex pattern arises from the combination of many admissible modes. The plot of the dispersion relation $\lambda$ vs the wavenumber $k^2$ obtained through the linear stability analysis is shown to the left of the corresponding predicted pattern. Red represents the highest $u$ concentration values and blue the lowest ones. To improve readability, when due to symmetry multiple degenerated modes exist for a given value of $k^2$, only one of them is shown. For example, mode $(1, 0, 1)$ in the legend also includes modes $(0, 1, 1)$ and $(1, 1, 0)$. The modes in the legend are listed in ascending value of their corresponding $k^2$.

### S5 Suppl. figures: Admissible modes change in a growing domain

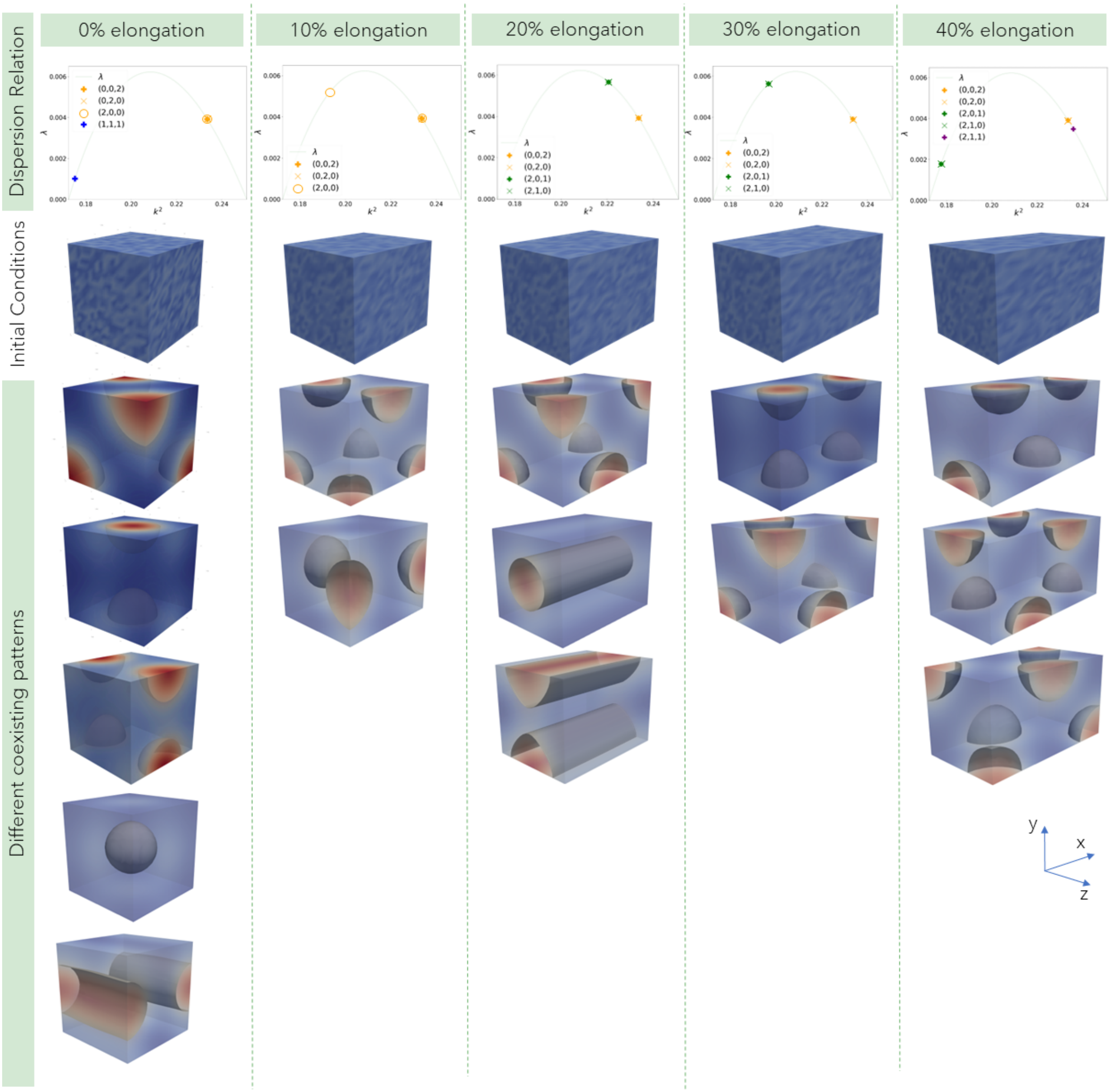


**Figure S3:** Admissible modes may change with different domain sizes, even when all other model parameters are the same. The plot of the dispersion relation $\lambda$ vs the wavenumber $k^2$ obtained through the linear stability analysis for each domain size of the results in figure 5 of the main text are shown. Several simulations for each domain size were performed, starting from different random initial conditions. Red represents the highest $u$ concentration values and blue the lowest ones.
